# HIV Protease Inhibitor Nelfinavir Targets Human DDI2 and Potentiates Proteasome Inhibitor-based Chemotherapy

**DOI:** 10.1101/2020.05.03.075572

**Authors:** Yuan Gu, Xin Wang, Yu Wang, Jie Li, Fa-Xing Yu

## Abstract

Proteasome inhibitors (PIs) are currently used in the clinic to treat cancers such as multiple myeloma (MM). However, cancer cells often rapidly develop drug resistance towards PIs due to a compensatory mechanism mediated by nuclear factor erythroid 2 like 1 (NFE2L1) and aspartic protease DNA damage inducible 1 homolog 2 (DDI2). Following DDI2-mediated cleavage, NFE2L1 is able to induce transcription of virtually all proteasome subunit genes. Under normal condition, cleaved NFE2L1 is constantly degraded by proteasome, whereas in the presence of PIs, it accumulates and induces proteasome synthesis which in turn promotes the development of drug resistance towards PIs. Here, we report that Nelfinavir (NFV), an HIV protease inhibitor, can inhibit DDI2 activity directly. Inhibition of DDI2 by NFV effectively blocks NFE2L1 proteolysis and potentiates cytotoxicity of PIs in cancer cells. Recent clinical evidence indicated that NFV can effectively delay the refractory period of MM patients treated with PI-based therapy. Our finding hence provides a specific molecular mechanism for combinatorial therapy using NFV and PIs for treating MM and probably additional cancers.

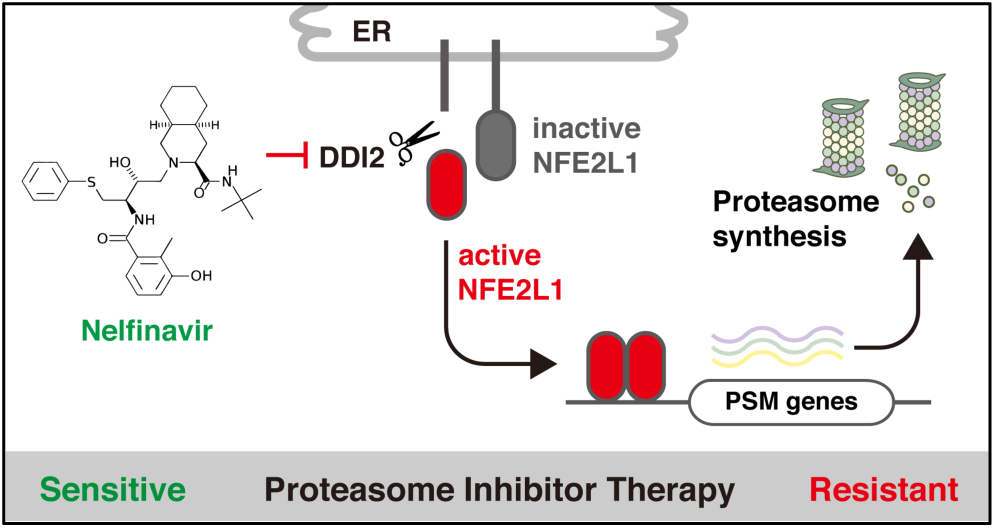

## Introduction

The ubiquitin-proteasome system (UPS) is a major protein degradation pathway in eukaryotic cells that participates in diverse biological and pathological processes, including immune defense, neuronal disorders, and tumorigenesis ^1-5^. In cancer cells, the proteasome is often overloaded with mutant or over-expressed proteins, making it a promising target for anti-cancer therapies ^3,6-8^. Several proteasome inhibitors (PIs) have been approved as first-line therapies for multiple myeloma (MM) and mantle cell lymphoma ^9,10^. Among them, Bortezomib (BTZ, marketed as Velcade) is a dipeptidyl boronic acid derivative that reversibly targets the active site of the β5-subunit of the 20S proteasome^11^. While PIs serve as the backbone therapy for MM, most patients relapse and become refractory towards PIs ^12-16^. In addition, despite a strong dependence on functional proteasome system, most cancer types, especially solid tumors, are resistant to PIs ^17,18^.

The development of resistance to PIs is most-likely dependent on compensatory proteasome synthesis. Nuclear Factor, Erythroid 2 Like 1 (NFE2L1, also known as NRF1 for NFE2-Related Factor 1), a CNC-type bZIP family transcription factor, serves as a master regulator for proteasome synthesis. In steady state, NFE2L1 translocates from the endoplasmic reticulum (ER) lumen to the cytoplasm by valosin containing protein (VCP/p97), and is quickly degraded by proteasome via the ER-associated degradation (ERAD) pathway ^19^. In the presence of PIs, cytosolic NFE2L1 is de-N-glycosylated by N-glycanase 1 (NGLY1) ^20,21^, and subsequently cleaved by aspartic protease DNA damage inducible 1 homolog 2 (DDI2) ^22^. The cleavage generates a C-terminal fragment of NFE2L1 which retains DNA-binding and transcription-activating properties. In the nuclei, the NFE2L1 C-terminal fragment induces expression of almost all the proteasome subunit genes ^23,24^. Newly synthesized proteasomes in turn dilute the efficacy of PIs and reestablish a balanced cellular protein turnover. This molecular strategy is known as the “bounce-back” mechanism for cells to handle stress upon proteasome inhibition ^19,25^ (Extended Data Fig.1a).

To reduce drug resistance towards PI-based chemotherapy, new proteasome synthesis should be minimized, and several enzymes in the “bounce-back” loop may serve as molecular targets. Very recently, it has been shown that NGLY1 inhibitors potentiate cytotoxicity of PIs ^20,21^, and inhibition of VCP/p97 induces cancer cell death ^26-28^. Moreover, DDI2 deficiency can attenuate the transcriptional activity of NFE2L1 and potentiate cytotoxicity of PIs ^29^. Compared to VCP/p97 and NGLY1, DDI2 is a more specific molecular target, because so far only NFE2L1 and its homolog NFE2L3 have been identified as substrates of DDI2. Hence, a small-molecule inhibitor targeting DDI2 has a great potential to improve PI-based chemotherapy.

In this report, we show that Nelfinavir (NFV), an approved drug for treating HIV infection, can directly inhibit DDI2. Moreover, NFV can effectively block accumulation of active NFE2L1 in the presence of PIs, and potentiate cytotoxicity of PIs in different cancer cells. The inhibition of DDI2 and NFE2L1 activation by NFV serves as a molecular mechanism underlying the synergistic effects of NFV and PI for combinatorial cancer therapy in different animal models and clinical trials.

## Results

### HIV protease inhibitor NFV represses DDI2-mediated NFE2L1 cleavage

Human DDI2 has a ubiquitin-like domain (UBL) at the amino-terminus (N) and a retroviral protease-like domain (RVP) domain near the carboxyl-terminus (C), and the RVP domain is responsible for its protease activity ^30^(Fig. 1a). The RVP domain is highly conserved in DDI2’s orthologs and retroviral proteases such as HIV protease ^30-32^. When spatially aligned, RVP domain structures of DDI2 and HIV protease largely overlapped, indicating that these two structures shared high degree of similarity (Fig. 1c). Hence, HIV protease inhibitors, many of which directly target the catalytic pocket of HIV protease, may also bind with RVP domain of DDI2 and affect DDI2 protease activity.

**Figure 1.**
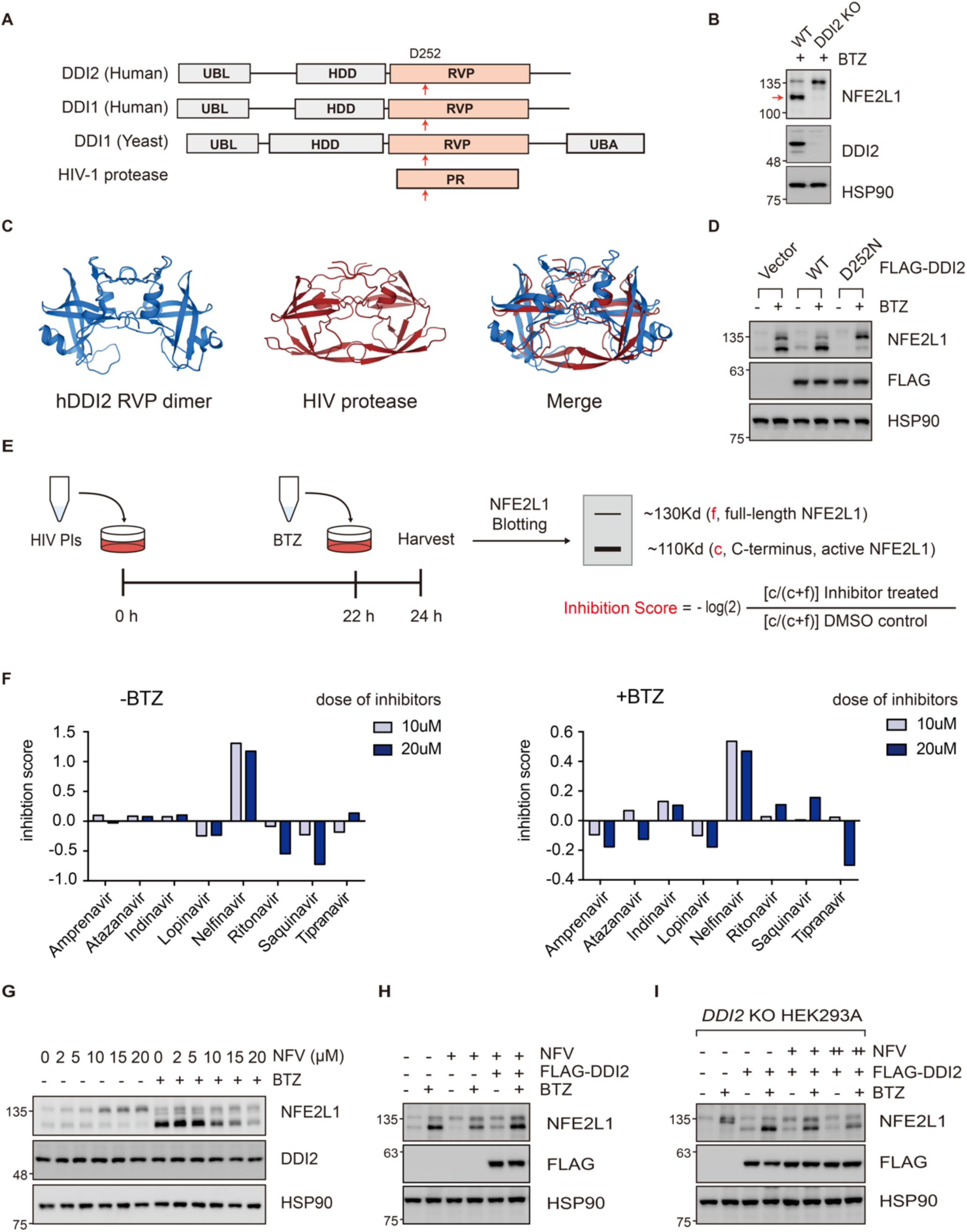
HIV protease inhibitor NFV represses DDI2-mediated NFE2L1 cleavage. (A) Domain organization of DDI2 homologs and HIV protease. UBL: ubiquitin-like domain; UBA: ubiquitin-associated domain; HDD: helical domain; RVP: retroviral protease-like domain; PR: protease. Arrow indicated the active aspartate. (B) Structures of human DDI2 RVP domain (blue) and HIV protease (red). (C) The cleavage of NFE2L1 was completely abolished in DDI2 KO cells, arrow indicated the C-terminal fragment of NFE2L1. (D) Overexpression of wildtype(WT) DDI2 promoted NFE2L1 processing while protease-dead DDI2 mutant (D252N) inhibited the processing. BTZ (100nM) was added 2 hours before harvest. (E) Schematic of small-scale screen for DDI2 inhibitors. (F) Relative inhibition of NFE2L1 cleavage by different small molecules. (G) NFV inhibited the processing of NFE2L1 in a dose-dependent manner. HEK293A were treated with different doses of NFV for 24 h. BTZ (100 nM) was added 2 h before cells were harvested. (H) DDI2 overexpression diluted the efficacy of NFV (10 µM, 24 h) on NFE2L1 cleavage. BTZ (100 nM) was added 2 h before cells were harvested. (I) NFV inhibited the activity of ectopically expressed DDI2 in DDI2 KO HEK293A cells. Cells were treated with 10 µM (+) or 20 µM (++) NFV for 24 h followed by BTZ (100 nM) for 2 h.

To test potential effects of HIV protease inhibitors on DDI2 activity, we utilized NFE2L1 cleavage as a reporter for DDI2 activity (Fig. 1e). The cleavage of NFE2L1 was a faithful readout of DDI2 activity, as it was completely abolished in *DDI2* knockout (KO) HEK293A cells and restored upon ectopic DDI2 expression (Fig. 1b, 1i). In the screen, cells were pre-treated with HIV protease inhibitors, including Amprenavir, Atazanavira, Indinavir, Lopinavir, NFV, Ritonavir, Saquinavir, or Tipranavir, followed by treatment with BTZ (or without), and the formation of active NFE2L1 by cleavage was monitored by immunoblotting (IB, Fig. 1f, Extended Data Fig. 2a). Among different HIV protease inhibitors, NFV inhibited NFE2L1 processing in a dose-dependent manner, with 20 µM NFV showing significant (∼75% in HEK293A cells) inhibition on NFE2L1 cleavage in various cell types including colorectal cancer cells and MM cell lines (Fig. 1g, Extended Data Fig. 3a-c). Moreover, the efficacy of NFV on NFE2L1 cleavage was largely diluted upon overexpression of DDI2 (Fig. 1h). Recently, NFE2L3, a homolog of NFE2L1, was reported as another DDI2 substrate ^33^, and the cleavage of NFE2L3 by DDI2 was also inhibited by NFV (Fig. Extended Data Fig. 3d). Together, these results indicate that NFV can effectively inhibit DDI2 activity.

### NFV directly targets human DDI2

To determine whether DDI2 is a direct target of NFV, we first conducted cellular thermal shift assay in which ligand-binding would increase a target protein’s thermodynamic stability over a range of temperatures ^34-36^. Cells were treated with or without NFV for 1 h, and intact cells in suspension were incubated at gradually increasing temperatures. The presence of NFV significantly increased DDI2 protein stability, indicating a direct binding between DDI2 and NFV (Fig. 2a). To gain structural insight of the interaction between NFV and DDI2, we performed computer-aided docking studies. Guided by the structure of NFV-bound HIV protease ^37^, we found that NFV could easily fit into the pocket of the DDI2 RVP domain dimer, with binding energies of the chosen clusters for NFV ranging from −9.38 to −9.46 kcal/mol (Fig. 2b). The central hydroxyl group of NFV bound to the catalytic aspartate of DDI2, similar to that of HIV protease (Fig. 2b). Hence, NFV most likely interacts with the catalytic center of DDI2 directly and inhibits DDI2 protease activity.

**Figure 2.**
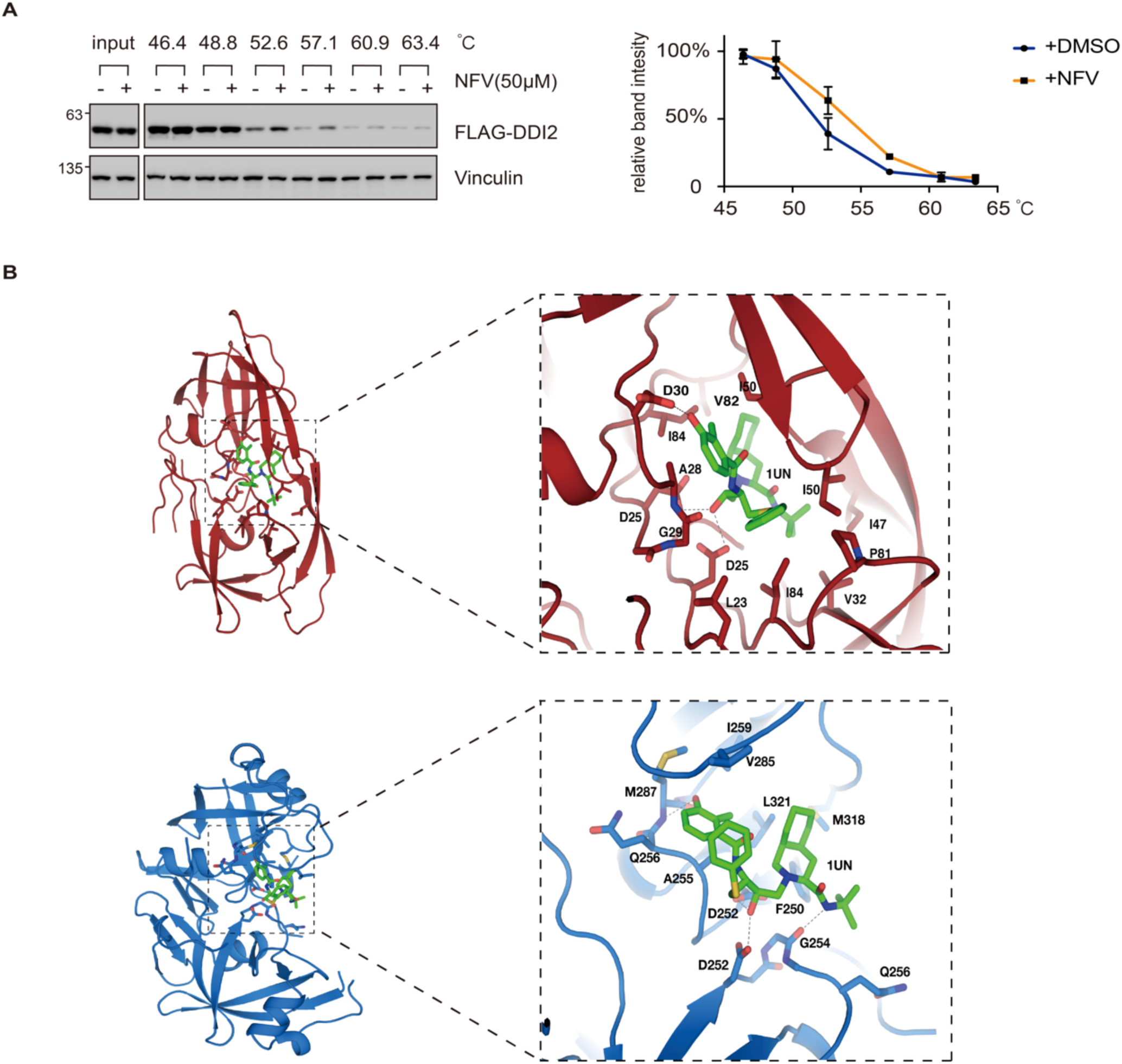
NFV directly targets human DDI2. (A) NFV stabilized DDI2 in a cellular thermal shift assay. HEK293A cells with or without NFV treatment were incubated at different temperatures, and DDI2 turnover was monitored by immunoblotting (left) and quantified (right, melting curve for results of three independent experiments). (B) Docking studies for possible conformations of NFV with human DDI2. Blue: human DDI2 RVP domain. Red: HIV protease. Green: NFV. Note the coordination between NFV with aspartic acids (D252 in DDI2).

DDI1 is a homolog of DDI2 and is mainly expressed in the testis (www.proteinatlas.org). DDI1 shares high sequence similarity with DDI2, suggesting a role in NFE2L1 processing and potential interaction with NFV (Extended Data Fig. 4a). Indeed, in *DDI2* KO cells, ectopically expressed DDI1 fully supported NFE2L1 cleavage. Moreover, NFV inhibited NFE2L1 cleavage in DDI1-reconstituted cells (Extended Data Fig. 4b-d). These results suggest that both DDI1and DDI2 are molecular targets of NFV.

### NFV blocks compensatory proteasome synthesis mediated by DDI2 and NFE2L1

It has been shown previously that DDI2 deficiency is able to switch off NFE2L1 processing and block new proteasome synthesis under proteasome inhibition ^22,29^. In the presence of BTZ, NFV effectively repressed active NFE2L1 accumulation, as well as the nuclear translocation of NFE2L1 (Fig. 3a-b). Active NFE2L1 in the nuclei worked as a transcription factor to increase the expression of genes encoding different proteasome subunits. As expected, NFV reduced the expression of proteasome subunit genes under BTZ treatment (Fig. 3c). Together, these lines of evidence indicate that, by inhibiting DDI2 activity and NFE2L1 activation, NFV is effective in disrupting the compensatory signaling (the “bounce-back” mechanism) under proteasome inhibition.

**Figure 3.**
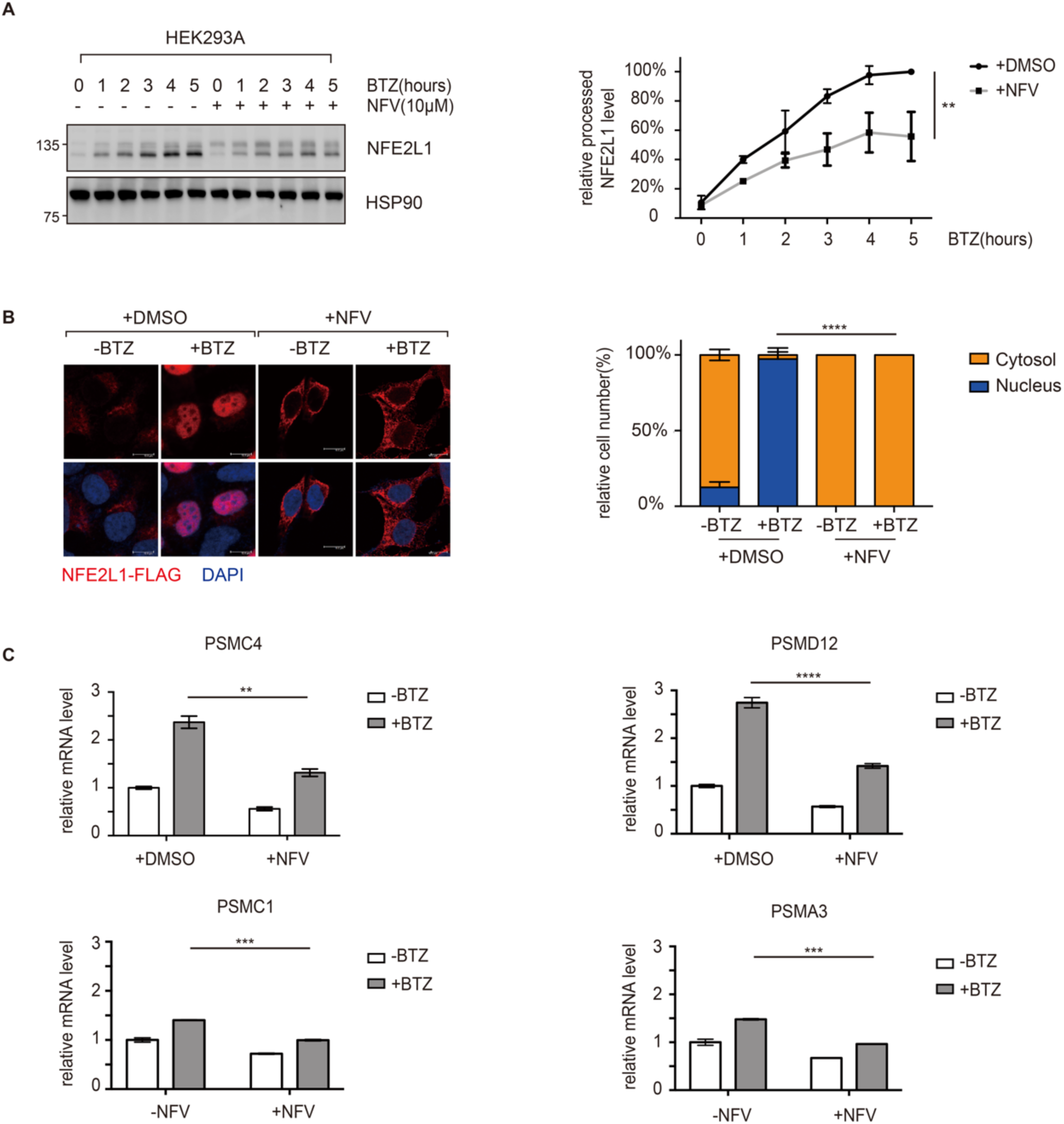
NFV blocks compensatory proteasome synthesis mediated by DDI2 and NFE2L1. (A) NFV delayed the accumulation of processed NFE2L1 under proteasome inhibition. HEK293A cells were treated with 10 µM NFV or DMSO for 24 h. BTZ (100 nM) was added for the indicated times before cells were harvested. Cleavage of NEF2L1 were determined by immunoblotting and quantified (right, summary of three independent experiments). (B) NFV blocked nuclear translocation of NFE2L1. HEK293A cell were transfected with NFE2L1 with FLAG tag at C-terminus, and subcellular localization was determined by immunofluorescence using FLAG antibody. Cells with cytoplasmic or nuclear NFE2L1-FLAG staining were counted (right). (C) The mRNA expression of proteasome subunit genes was repressed by NFV. The mRNA levels of HCT116 cells with or without NFV treatment (10 µM, 24 h) or BTZ treatment (100 nm, 12 h) were determined by quantitative PCR, and normalized by GAPDH mRNA levels. Mean and standard error were presented (*p<0.05, **p < 0.005, ***p < 0.0005, ****p <0.00005, ns = not significant; n=3; t test).

### NFV promotes cancer cell death induced by PI

Since NFV is effective in inhibiting compensatory proteasome synthesis, we next tested if NFV can improve cytotoxicity of PIs towards cancer cells. As determined by CCK8 assays, BTZ alone showed a lethal dose (LD_50_) at 10 nM for HCT116 colorectal cancer cells (Fig. 4a-b). Overexpressing DDI2-D252N, a protease-dead DDI2 mutant, functioned in a dominant negative way (Fig. 1d) and significantly decreased LD_50_ of BTZ (Fig. 4b). This result confirmed that targeting DDI2 could increase sensitivity of cancer cells to PIs. In line with data from DDI2-D252N overexpression, NFV treatment also decreased LD_50_ of BTZ. Interestingly and in contrast, DDI2 overexpression significantly increased LD_50_ of BTZ, and NFV reversed this effect (Fig. 4b), consistent with their effects on NFE2L1 processing (Fig 1i). Hence, high DDI2 activity is protective for cells under proteasome stress, and targeted inhibition of DDI2 by NFV can sensitize cancer cells to death. Meanwhile, NFV treatment showed no further effect on the LD_50_ of BTZ in DDI2(D252N)-overexpressing HCT116 cells (Fig 4b), indicating that NFV’s effect on PI sensitivity was mediated by DDI2 inhibition.

**Figure 4.**
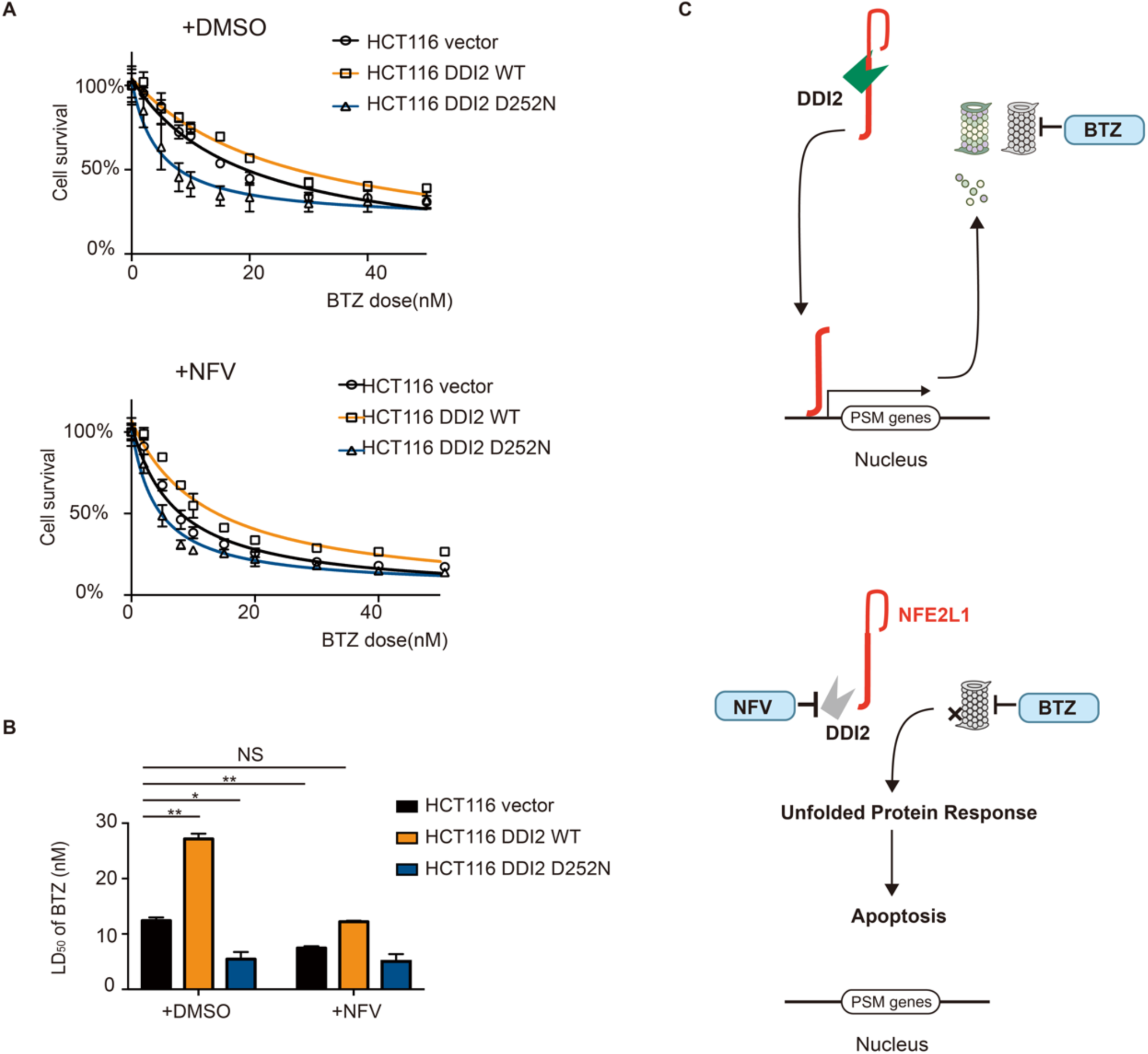
NFV promotes cancer cell death induced by PI. (A) NFV and BTZ synergistically promoted cell death. Control, DDI2-overexpressing, or DDI2 D252N-expressing cells were treated with different doses of BTZ together with DMSO or 10 µM of NFV for 24 h, survived cells were determined by CCK8 assay. (B) LD_50_ of different groups were calculated according the survival curve in (A). Mean and standard error were presented (*p<0.05, **p < 0.005, ns: not significant, n=3, t test). (C) A proposed model of NFV and PI (BTZ) combinatorial therapy.

### Clinical evidence of NFV in PIs-refractory MM Treatment

NFV is currently used in the clinic as a therapy for HIV infection ^38^. Based on our findings, NFV may be repurposed as an anti-cancer agent, especially when used in combination with PIs. Recently, multiple clinical studies indicate that the combination of NFV and BTZ had a great response in the refractory period of MM patients ^39-41^ (Table 1), even though a clear molecular mechanism was lacking. Hence, the direct inhibition of DDI2 by NFV, and the resulting repression of NFE2L1 activity and proteasome synthesis, represents a robust mechanism underlying the synergestic effect between NFV and PIs in chemotherapy.

**Table 1.** Clinical evidence of NFV in PIs-refractory MM Treatment.

| TRIAL IDENTIFIER | PHASE | STATUS | STUDY TITLE | CONDITIONS | INTERVENTION | RESPONSE | REFERENCE |
| --- | --- | --- | --- | --- | --- | --- | --- |
| NCT01164709 | 1 | Completed | Nelfinavir Mesylate and Bortezomib in Treating Patients with Relapsed or Progressive Advanced Hematologic Cancer | Leukemia, Lymphoma, Mature T-cell and NK-cell Neoplasms and Multiple Myeloma and Plasma Cell Neoplasm; Relapsed or Progressive | NFV+ BTZ | 83% ORR, 50% PR, 33% MR* | Driessen et al., 2016 |
| NCT02188537 | 2 | Completed | Nelfinavir as Bortezomib-sensitizing Drug in Patients with Proteasome Inhibitor-nonresponsive Myeloma | Myeloma; Proteasome Inhibitor-nonresponsive | NFV + BTZ + dexamethasone | 74% ORR, 15% VGPR, 50% PR, 9% MR | Driessen et al., 2018 |
| NCT01555281 | 1/2 | Active, not recruiting | Nelfinavir and Lenalidomide/Dexamethasone in Progressive Multiple Myeloma | Multiple Myeloma; lenalidomide-refractory | NFV + Lenalidomide+ dexamethasone | 55% ORR, 10% VGPR, 21% PR, 24% MR | Hitz et al., 2019 |

\*Extension cohort with 6 patients. MM, multiple myeloma; MR, minimal remission; ORR, objective response rate; PI, proteasome inhibitor; PR, partial response; VGPR, very good partial response.

## Discussion

In summary, we have identified NFV as the first small-molecule inhibitor of human DDI2 with a mechanism that likely involves direct interaction with the catalytic center of DDI2. The inhibition of DDI2 by NFV blocks cleavage and activation of NFE2L1 upon proteasome inhibition, and represses compensatory proteasome synthesis. Insufficient proteasome activity may cause accumulation of defective proteins and lead to cell death (Fig. 4c). Indeed, NFV and PIs have shown synergistic effects in killing cancer cells *in vitro* and overcoming drug resistance in refractory period of MM patients clinically. Hence, our study provides a novel molecular mechanism for the clinical efficacy of NFV and BTZ combinatorial cancer therapy.

Our results indicate that NFV may serve as a lead compound for the development of more potent inhibitors of DDI2. However, our *in silico* modeling of the interaction between NFV and the catalytic center of DDI2 is mainly based on the structural information of HIV protease bound with NFV. Whereas, among different HIV protease inhibitors tested, only NFV showed potent inhibitory effect on DDI2 activity, suggesting some differnces between DDI2 and HIV protease. In the future, a crystal strucute of DDI2 bound with NFV would give more insights regarding to the refinement of DDI2 inhibitors.

The clinical effect of NFV in conjunction with BTZ is currently tested on MM patients. Our work indicates that DDI2 activity serves as a protective factor for cancer cells under proteasome stress, so cancers with high DDI2 expression, such as Esophageal Carcinoma, Acute Myeloid Leukemia, and Esophageal Carcinoma, may also be sensitive to NFV and PI combinatorial therapy (Extended Data Fig. 5a). Hence, the NFV and BTZ combinatorial therapy may also effective in treating some solid tumors with high DDI2 activity.

## Experimental procedures

### DNA constructs

Human NFE2L1 and DDI2 cDNAs were cloned using total RNA extracted from HEK293A cells. Primers used were shown in extended data table 1. Human DDI1 cDNA was synthesized by Genewiz, Suzhou, China. PCR was performed using PrimeSTAR Max DNA polymerase (Takara Bio, Shiga, Japan). Amplified fragments were subcloned into pLVX lentiviral vector and all plasmids were confirmed by Sanger sequencing.

### Cell culture and transfection

HEK293A, HEK293T, HCT116 and HCT15 cells were cultured in DMEM (Corning) containing 5% FBS (Gibco) and 50 µg/mL penicillin/streptomycin (P/S). RPMI8226 cells were cultured in RPMI1640 (Corning) containing 10% FBS (Gibco) and 50 µg/mL penicillin/streptomycin (P/S). All cell lines were maintained at 37°C with 5% CO2. Cells were transfected with plasmid DNA using PolyJet DNA In Vitro Transfection Reagent (Signagen Laboratories, Gaithersburg, USA) according to manufacturer’s instructions.

### Chemicals

The following chemicals were used in this study. HIV protease inhibitors including Amprenavir, Atazanavira, Indinavir, Lopinavir, NFV, Ritonavir, Saquinavir, Tipranavir and proteasome inhibitor BTZ were either obtained from the Selleck protease inhibitor kit or purchased separately. Large package of NFV Mesylate was purchased from MCE.

### Immunoblotting

Immunoblotting was performed using standard protocol. The following primary antibodies were used in Immunoblotting: Vinculin (CST, E1E9V), NFE2L1(CST, D5B10), FLAG-HRP (Sigma, A8592), HSP90(BD, 610418). Vinculin and NFE2L1 were diluted 1:1000 while the other two were diluted 1:10000 in TBST containing 5% BSA. Data were quantified using Image J software.

### Immunofluorescence

HEK293A cells stably overexpressing NFE2L1-flag were seeded on coverslips. After treatment, cells were fixed with 4% paraformaldehyde-PBS for 10 min and permeabilized with 0.1% Triton X-100 in TBS. After blocking in 3% BSA and 3% goat serum in PBS for 1 h, cells were incubated with FLAG antibody (CST, D6W5B) overnight at 4°C. After three washes with PBS, cells were incubated with Alexa Fluor 488- or 555-conjugated secondary antibodies (Invitrogen, 1:1000 diluted) for 1 h at room temperature. Slides were then washed three times and mounted. Images were captured using Olympus confocal microscopy.

### Cellular Thermal Shift Assay

Cellular thermal shift assay was conducted according to the protocol as previously described^34^. HEK293A cells stably expressing FLAG-DDI2 were treated with 50 µM NFV or DMSO for 1 h, and cells were collected and washed with PBS buffer three times to avoid excess compound residue. Cells in suspension were equally dispensed into 0.2 ml PCR tubes (3 million cells per tube), incubated at preset temperatures for 3 min on a PCR instrument, and freeze-thawed twice using liquid nitrogen. Samples were centrifuged and the supernatants were analyzed by immunoblotting. All experiments were performed in triplicates.

### Docking studies for possible conformations of NFV with human DDI2

To gain structural understanding and visualization of the interaction between NFV and hDDI2, docking studies were performed using Autodock4^42^. The hDDI2 structure was prepared using previously determined structure of DDI2 (from Gln232 to Glu328, PDB No. 4RGH). All bound waters were removed and then added for hydrogens. All partial atomic charges were assigned automatically using AutoDockTools (ADT). The coordinates of NFV were generated from the structure of HIV-1 protease co-crystallized with NFV (PDB No. 1OHR). The hydrogen atoms and Gasteiger charges were then assigned to the ligand using ADT. The interaction was modeled with the Lamarckian genetic algorithm. The clusters with lower energies and reasonable conformations were chosen as solution. The binding energies of the chosen clusters for NFV ranged from −9.38 to −9.46 kcal/mol. The orientation of NFV was further confirmed using LeDock (data not shown)^43^.

### Quantitative RT-PCR

Total RNA was extracted using the Takara MiniBEST universal RNA Extraction kit (Takara Bio, Shiga, Japan). cDNA was generated using the TransScript First-Strand cDNA synthesis kit (TransGen Biotech, Beijing, China), and quantitative qPCR was conducted using SYBR Green qPCR Master Mix (Takara Bio, Shiga, Japan) on a 7500 Real-Time PCR systems (Applied Biosystems). Relative abundance of mRNA was calculated by normalization to GAPDH mRNA. Primers used are listed in extended data table 2.

### CCK8 cytotoxic assay

CCK8 cytotoxic assay was conducted by CCK-8 Cell Counting Kit (Yeasen, Shanghai, China) according to manufacturer’s instructions. 10,000 HCT116 cells were seeded on 96-well plates. Cell were treated with different doses of drug for 24 h before analysis, and DMSO was used as control.

## ASSCIATED CONTENT

### Supporting Information

Supplementary tables and figures are included in the end of this PDF

## AUTHOR INFORMATION

### Author Contributions

G. Y., and F.X. Y. designed the experiments, analyzed data, and wrote the manuscript. G. Y., X. W., Y. W., and J. Li performed experiments and analyzed data.

## ACKNOWLEDGMENT

This study is supported by grants from the National Natural Science Foundation of China (81772965), the National Key R&D program of China (2018YFA0800304), Science and Technology Commission of Shanghai Municipality (19JC1411100), and Shanghai Municipal Commission of Health and Family Planning (2017BR018) to FXY.

## ABBREVIATIONS

DDI2: DNA damage inducible 1 homolog 2;
NFE2L1: nuclear factor erythroid 2 like 1;
NFE2L3: nuclear factor erythroid 2 like 3;
NGLY1: N-glycanase 1;
P97/VCP: valosin containing protein.

---

**Extended Data Table 1.**
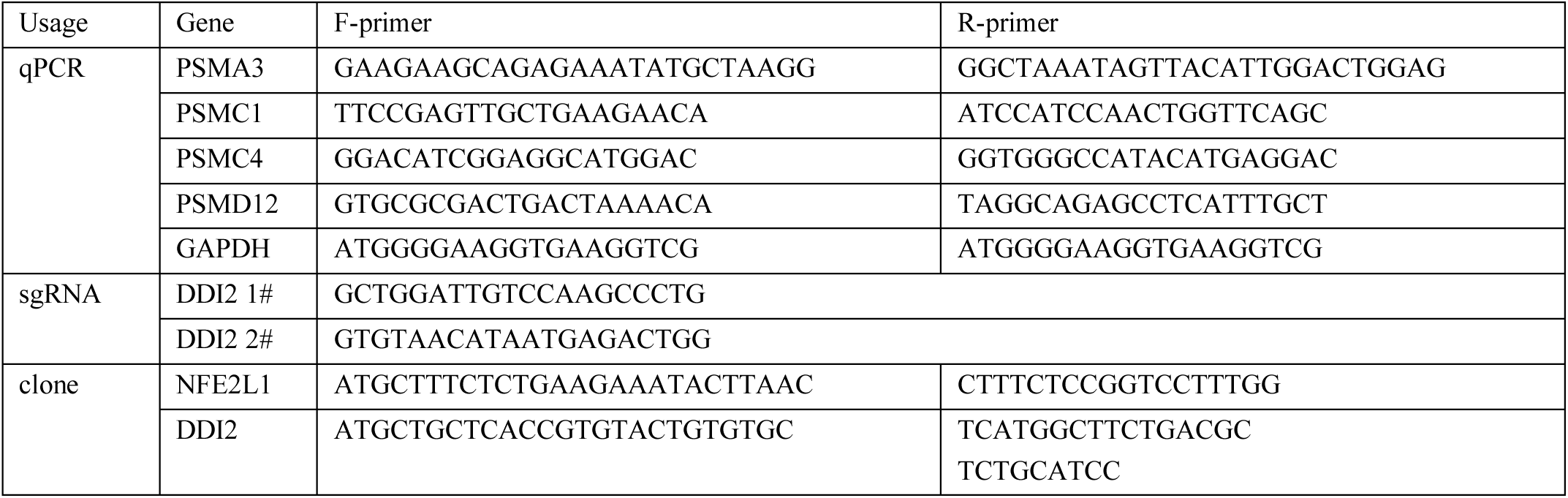
Oligos used in this study.

**Extended Data Figure 1.**
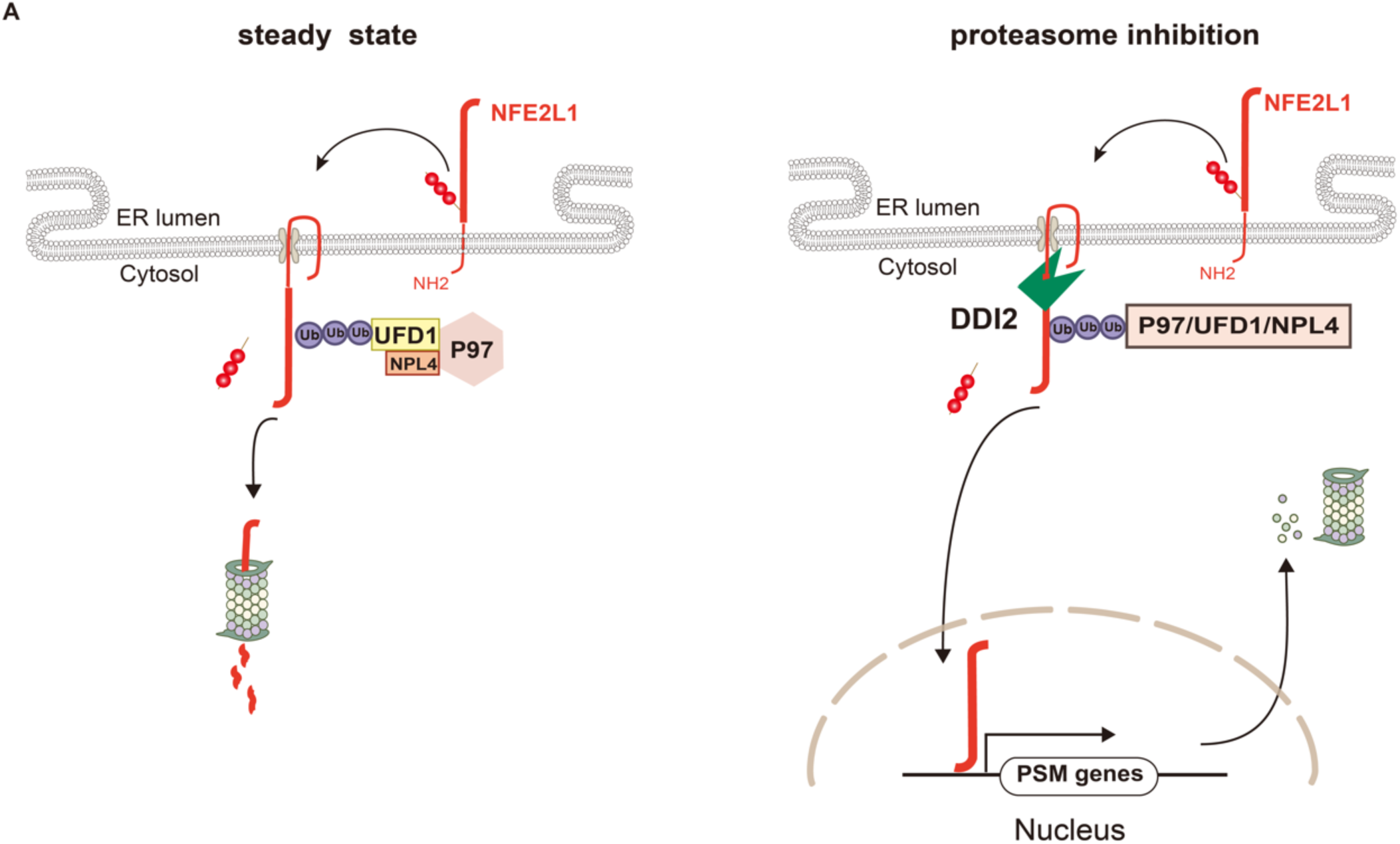
Activation of NFE2L1 and compensatory proteasome synthesis. UFD1: Ubiquitin recognition factor in ER-associated degradation protein 1; NPL4: Nuclear protein localization protein 4 homolog; Ub: ubiquitin; PSM genes: proteasome subunit genes.

**Extended Data Figure. 2.**
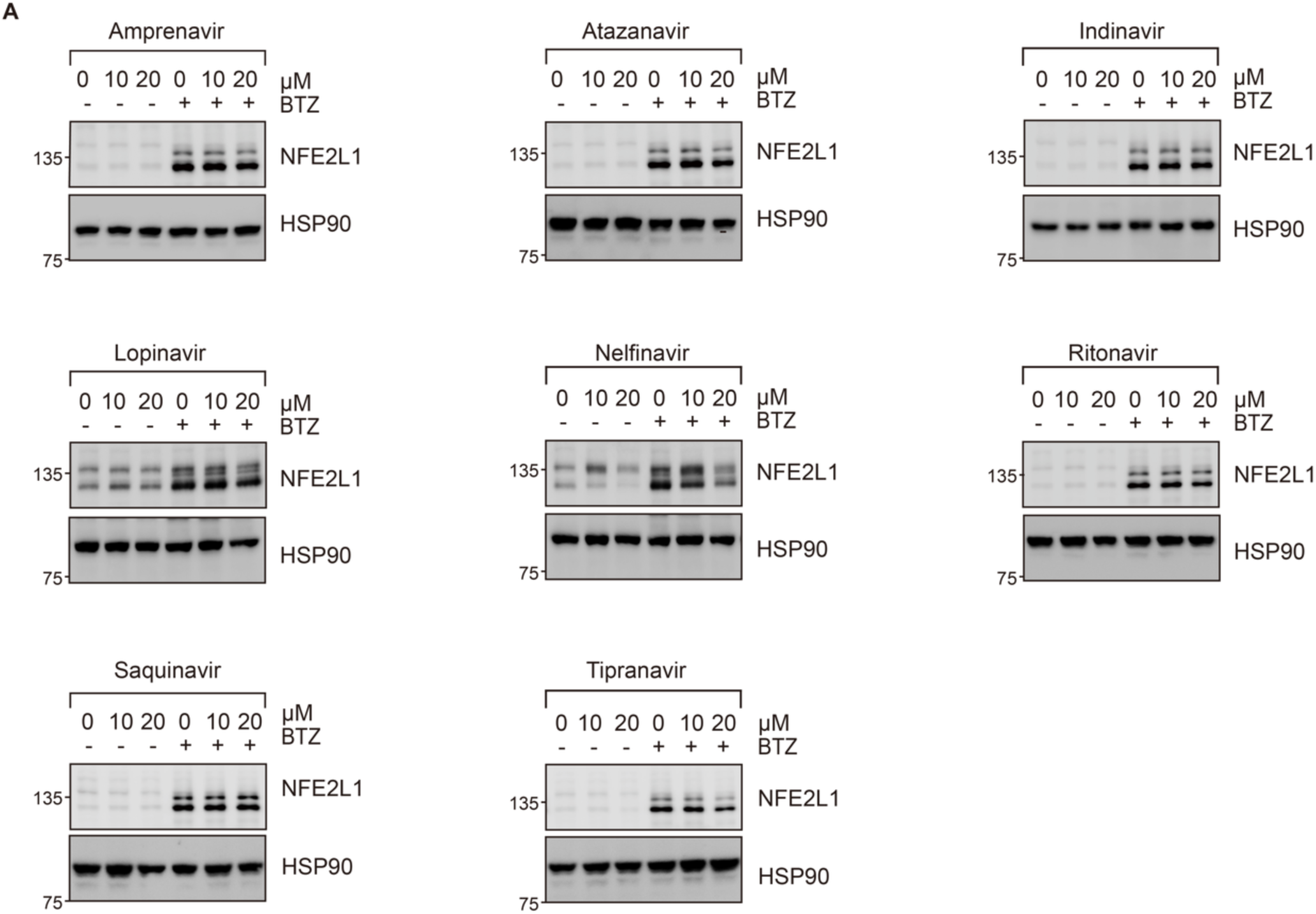
Representative immunoblotting results used in Fig 1C.

**Extended Data Figure. 3.**
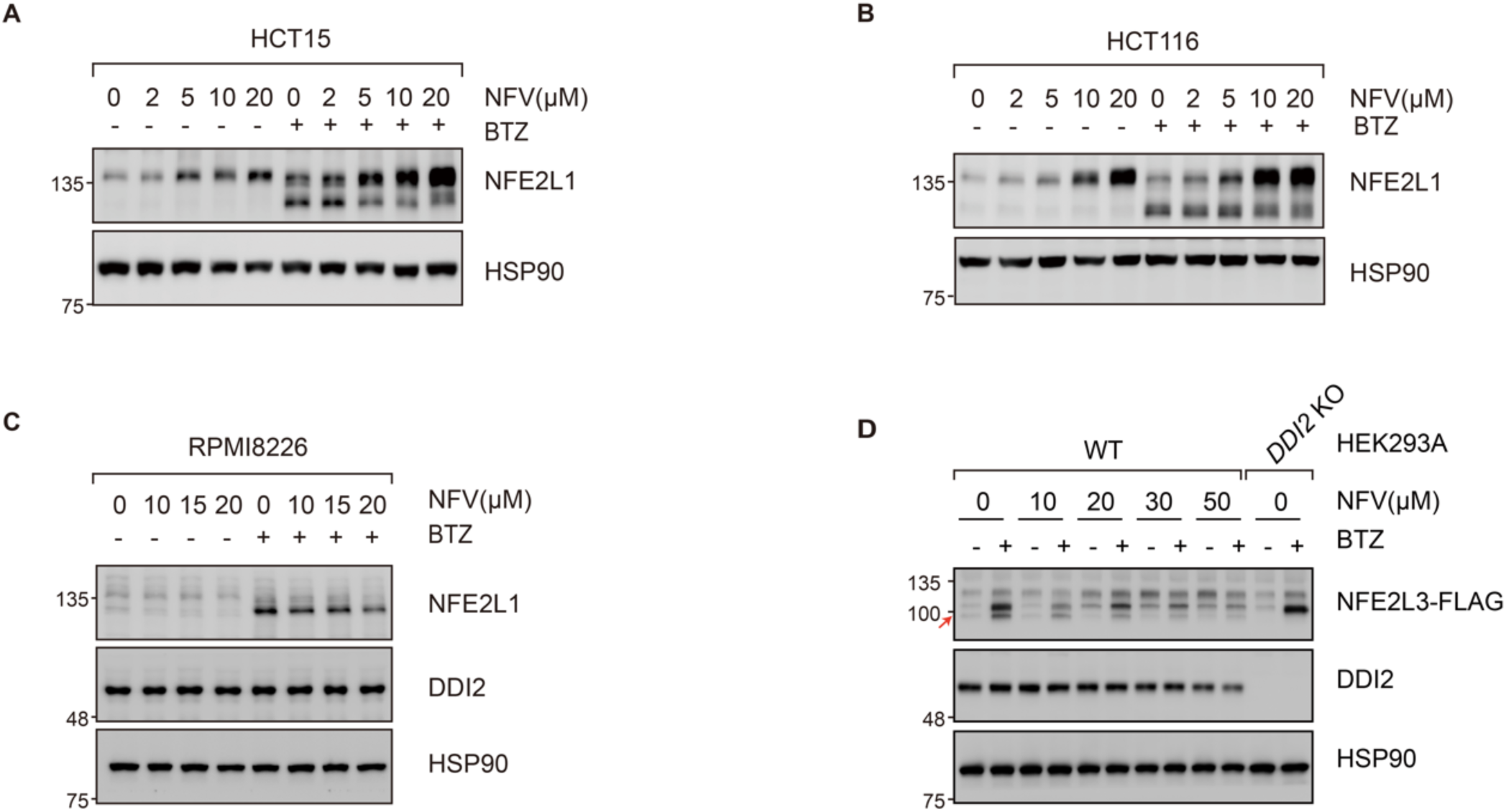
NFV inhibits NFE2L1 activation in different cancer cells, as well as NFE2L3 processing in HEK293A cells. NFV inhibited NFE2L1 cleavage upon proteasome inhibition in HCT15 (A), HCT116 (B) and RPMI8226 (C) cells. Cells were treated with different doses of NFV for 24 h. BTZ (100 nM) was added for 2 h before cells were harvested. (D) NFV inhibited the processing of NFE2L3. HEK293A overexpressing NFE2L3-FLAG were treated with different doses of NFV for 24 h followed by BTZ treatment (100 nM) for 2 h, arrow indicated the C-terminal fragment of NFE2L3.

**Extended Data Figure. 4.**
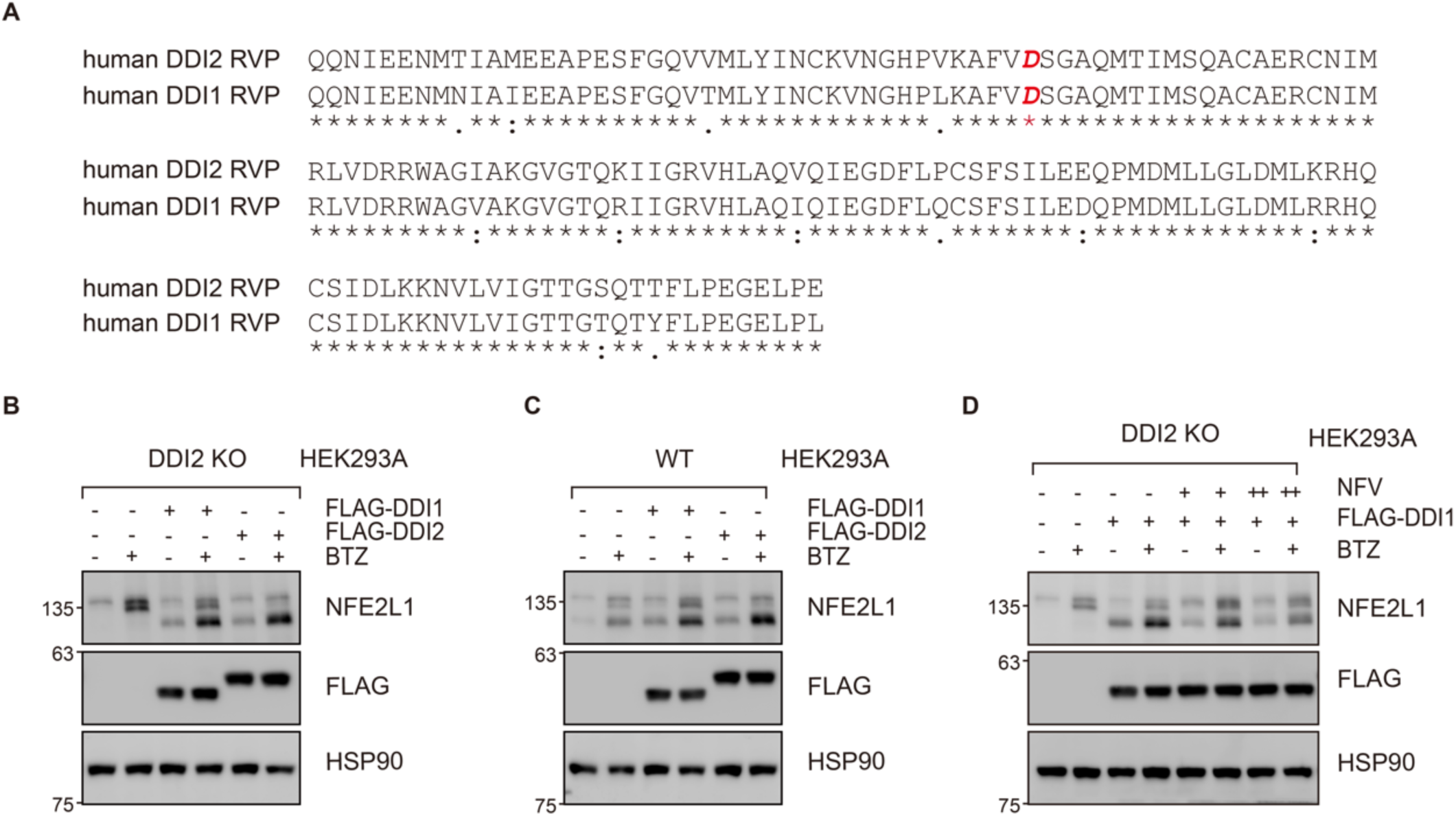
NFV targets human DDI1. (A) Amino acid sequence alignment of DDI1 and DDI2 RVP domain. (B) Ectopic expression of DDI1 in DDI2 KO HEK293A cells restored NFE2L1 cleavage. (C) DDI1 promoted NFE2L1 cleavage as DDI2 did. (D) NFV inhibited the activity of ectopic DDI1 in DDI2 KO HEK293A cells. Cells were treated with 10 µM (+) or 20 µM (++) NFV for 24 h. BTZ (100 nM) was added 2 h before cells were harvested.

**Extended Data Figure 5.**
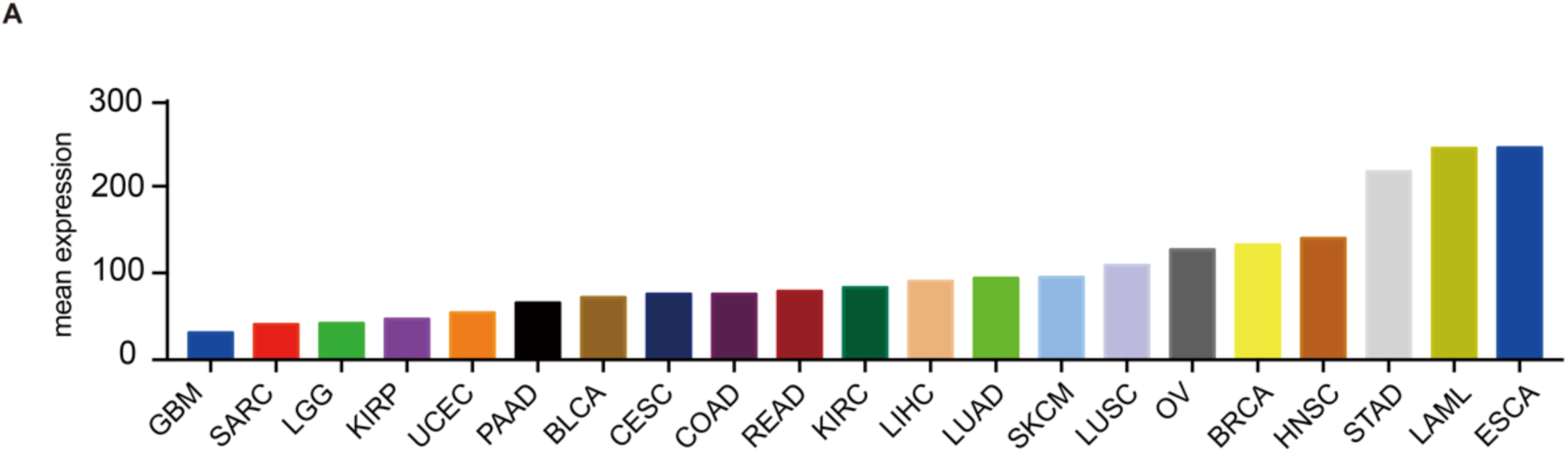
(A) Mean DDI2 mRNA level in different cancer types. Data from www.oncolnc.org. BLCA: Bladder Urothelial Carcinoma; BRCA: Breast invasive carcinoma; CESC: Cervical squamous cell carcinoma and endocervical adenocarcinoma; COAD: Colon adenocarcinoma; ESCA: Esophageal carcinoma; GBM: Glioblastoma multiforme; HNSC: Head and Neck squamous cell carcinoma; KIRC: Kidney renal clear cell carcinoma; KIRP: Kidney renal papillary cell carcinoma; LAML: Acute Myeloid Leukemia; LGG: Brain Lower Grade Glioma; LIHC: Liver hepatocellular carcinoma; LUAD: Lung adenocarcinoma; LUSC: Lung squamous cell carcinoma; OV: Ovarian serous cystadenocarcinoma; PAAD: Pancreatic adenocarcinoma; READ: Rectum adenocarcinoma; SARC: Sarcoma; SKCM: Skin Cutaneous Melanoma; STAD: Stomach adenocarcinoma; UCEC: Uterine Corpus Endometrial Carcinoma.

